# Integrated analysis of Tn-Seq and pangenome reveals core and lineage-specific essential genes in *Mycobacterium avium* subsp. *hominissuis*

**DOI:** 10.1101/2025.10.13.682076

**Authors:** Kotaro Sawai, Marie Ikai, Motoko Shinohara, Yukiko Nishiuchi, So Fujiyoshi, Yohei Doi, Tomotada Iwamoto, Kentaro Arikawa, Fumito Maruyama, Yusuke Minato

**Author notes:** Address correspondence to Yusuke Minato.

## Abstract

Pulmonary disease caused by nontuberculous mycobacteria (NTM-PD) is an emerging global health concern. Among NTM, *Mycobacterium avium* subsp. *hominissuis* (MAH) is the major causative agent of NTM-PD. Similar to *Mycobacterium tuberculosis* (*Mtb*), MAH exhibits lineage-specific geographical distributions and host adaptations. Here, we characterized three MAH strains from the residential bathrooms of MAH-PD patients in Japan. A genetic population clustering analysis revealed that the three strains belong to the East Asia (EA) lineages that are predominant in Japan and Korea. Pan-genome analysis using the publicly available complete genome sequences of MAH and the newly sequenced MAH strains identified 3,313 core genes that are conserved among distinct MAH lineages. Identification of essential genes in the three strains was conducted using transposon insertion sequencing (Tn-Seq), and their gene essentiality profiles were compared to those of a previously studied SC3 lineage strain, MAC109. Despite their genetic diversity, nearly all essential genes were derived from the core gene set. In addition, we identified a set of common essential genes for the EA and SC3 lineages, as well as lineage-specific essential genes. Our results highlight the evolutionary and clinical importance of lineage-specific adaptations in MAH.

**Importance:** By integrating transposon insertion sequencing with pan-genome analysis, we provide the first systematic comparison of essential genes across multiple *Mycobacterium avium* subsp. *hominissuis* (MAH) strains. Although MAH strains exhibit remarkable genetic diversity, we found that MAH essential genes are primarily confined to the core genome of MAH. This essential plasticity highlights the evolutionary strategies that underpin MAH survival across diverse environments and patient populations. Recognizing this interplay provides a foundation for identifying robust drug targets and developing lineage-informed therapies for MAH infection.

## Introduction

Over the past two decades, pulmonary diseases caused by nontuberculous mycobacteria (NTM-PD), a diverse group of environmental opportunistic pathogens, have been increasingly diagnosed and reported worldwide (1, 2). The incidence of NTM-PD has risen steadily, with particularly high prevalence in East Asia. Among NTM-PD, *Mycobacterium avium* complex (MAC) pulmonary disease (MAC-PD) is the most common (2, 3). For example, MAC-PD accounts for about 90% of reported NTM-PD cases, and the incidence rate is estimated at 14.7 per 100,000 population in Japan (4).

*M. avium* subsp. *hominissuis* (MAH), the major causative agent of MAC-PD, is ubiquitous in natural and environmental sources, including soil, dust, and water systems (5, 6). Recent advances in comparative genomics have highlighted the remarkable genetic diversity of MAH isolates from both clinical and environmental sources (7–9). Based on phylogenetic and population structure analysis of MAH genomes, MAH has been divided into seven major lineages, including MAHEastAsia1 (EA1), MAHEastAsia2 (EA2), and sequence clusters (SCs) 1 to 5 (7–9). Similar to *M. tuberculosis* (*Mtb*), MAH exhibits lineage-specific geographical distributions and host adaptations, with the EA lineages being predominant in MAC-PD patients in Japan and Korea (10).

Essential genes, defined as genes indispensable for growth and/or survival, represent promising targets for the development of novel antimicrobial therapies. Transposon insertion sequencing (Tn-Seq) has emerged as a powerful tool to identify essential genes on a genome-wide scale (11–24). Recently, Tn-Seq has been applied to identify strain/lineage-specific essential genes (13, 16). However, the application of Tn-Seq in MAH has been limited to only two strains belonging to SC3 due to the difficulty of constructing a high-density transposon mutant library (22, 23). Consequently, essential genes in non-SC3 MAH strains remain poorly understood.

Here, we sequenced the genomes of three MAH strains isolated from the residential environments of MAC-PD patients in Japan and revealed that they belong to the EA1 and EA2 lineages. We generated highly saturated transposon mutant libraries from the three strains and performed genome-wide essentiality analysis using Tn-Seq. By comparing these data with previously published SC3 lineage data, we identified potential lineage-specific and core-essential genes, providing new insights into MAH biology and potential drug targets.

## Results

### Lineage classification and pan-genome architecture of MAH strains

We previously isolated MAH strains from the residential bathrooms of MAH-PD patients in Japan (25). Here, we determined the complete genome sequences of the three strains, OCU682, OCU683, and OCU803 (Table S1). The total genome sizes of the three strains ranged from 5.1 to 5.2 Mbp. Each strain harbored one to three plasmids, with plasmid sizes ranging from 17 to 151 kb. The number of predicted coding sequences (CDSs) ranged from 4,864 to 5,016.

To determine the lineage of the three strains, we performed a genetic population clustering analysis. We searched for available MAH genome sequences from the NCBI RefSeq database and obtained 182 genome sequences. Based on 41,005 core-genome SNPs, the 185 strains were classified into six lineages (Fig. 1, Table S2), consistent with previous reports (7–9).

**Figure 1.**
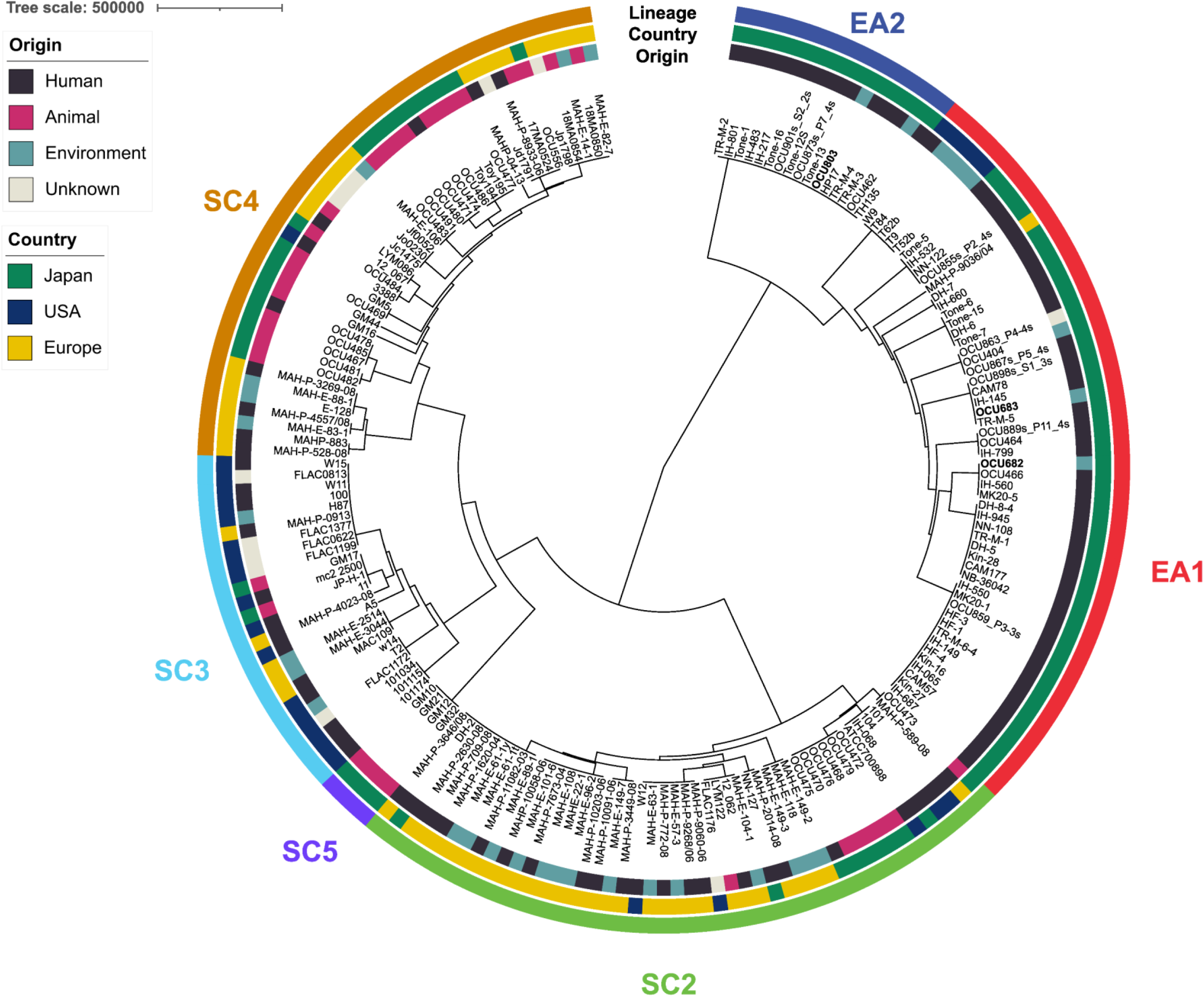
Population structure of *Mycobacterium avium* subsp. *hominissuis* (MAH) Complete linkage clustering of 185 isolates based on SNP distances. The tree scale (branch height) represents the increase in within-cluster variance at each node under Ward’s D2 criterion. The inner ring denotes the source of isolates: black for human, magenta for animal, turquoise for environment, and gray for unknown. The middle ring indicates the country of origin: green for Japan, navy for the USA, and yellow for Europe. The outer ring shows lineage classifications assigned by fastBAPS: green for SC2, light blue for SC3, brown for SC4, purple for SC5, red for EA1, and navy for EA2. Bold labels highlight isolates sequenced in this study.

Previously characterized 135 strains were classified into the same lineages, except for one strain, the A5 strain, which clustered into SC3 instead of SC1 (8, 9) (Table S2). The most well-characterized MAH standard strain, the MAH104 strain, was classified into the SC2 lineage.

The newly analyzed 50 strains were classified into five lineages: EA1, EA2, SC2, SC3, and SC4. OCU682 and OCU683 belonged to the EA1 lineage, while OCU803 was assigned to EA2 (Fig. 1, Table S2). The MAC109 and MAH11 strains, which were previously used to identify essential genes by Tn-Seq, were both classified into the SC3 lineage. Consistent with previous studies, strains classified into the EA1 and EA2 lineages were predominantly derived from human-associated samples in Japan, while strains classified into the SC2, SC3, and SC4 lineages were derived from various sources and regions (7–9).

### Core and accessory MAH genomes

We next aimed to understand how many genes are conserved among different lineages. To this end, 23 complete genome sequences of MAH were obtained from the NCBI RefSeq database. We also determined the complete sequences of OCU682, OCU683, and OCU803 strains. The pan-genome analysis of the 26 strains identified a total of 8,241 unique gene clusters. Of these, 3,313 were classified as core genes (conserved in ≥99% strains) (Figure 2A, Supplementary figure 1). A core gene-based phylogenetic tree revealed that strains from the EA1 and EA2 lineages clustered closely, while those from SC3 formed a distinct clade. The number of shell/soft-core (conserved in 15-99% strains) and cloud genes (conserved in 0-15% strains) varied across lineages (Figure 2B). Since each isolate contains a similar number of unique genes, the total accessory gene count increases with the number of isolates within a lineage (Figure S1).

**Figure 2.**
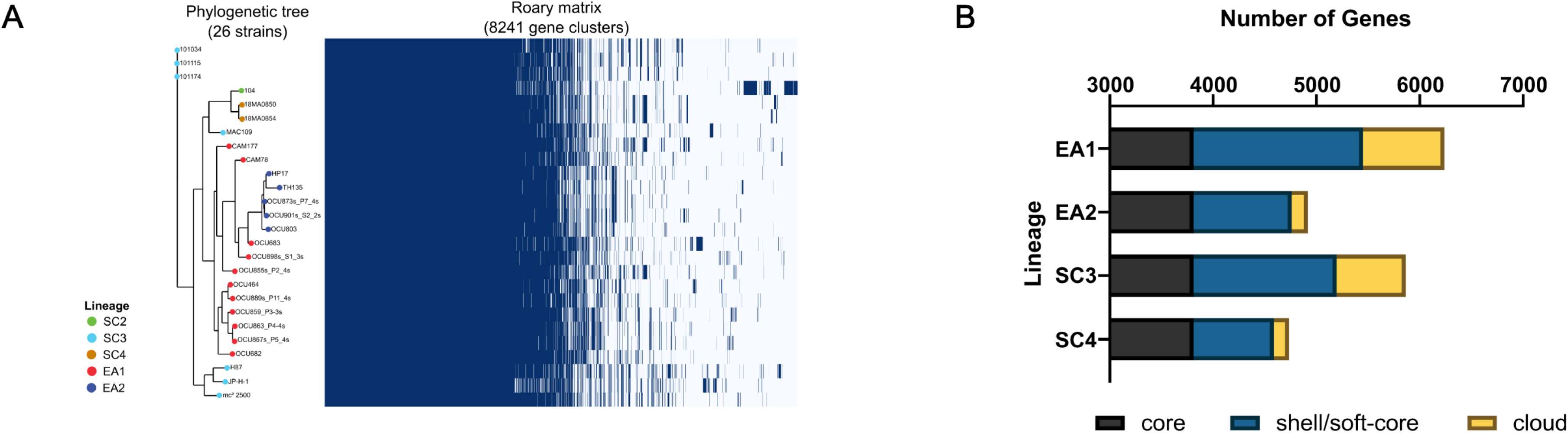
Pangenome structure and gene content distribution across 26 MAH isolates. A) The Roary matrix analysis identified 3,313 genes present in ≥99 % strains. The phylogenetic tree on the left was constructed using IQ-TREE with 1,000 bootstrap replicates. B) Each bar chart represents the number of core, soft-core/shell, and cloud genes within each lineage. Core genes: conserved in ≥99 % strains, soft-core genes: conserved in 95-99 % strains, shell genes: conserved in 15-95% strains, cloud genes: conserved in 0-15 % strains.

### Gene classification of three strains using TRANSIT

Because MAH strains generally show resistance to transduction by mycobacteriophage, generating a high-density transposon mutant library of MAH has been challenging (23). To overcome this challenge, we optimized the protocol for each strain by using distinct phage titers and the length of phage transduction time. As a result, we successfully obtained high-density transposon mutant libraries for all three strains.

Genome sequencing showed that OCU682, OCU683, and OCU803 contain up to 60,667 TA dinucleotides, with an additional up to 2,536 TA sites in their plasmids. To generate saturated transposon insertion mutant libraries from each strain, we collected at least 750,000 colonies of transposon mutants from each replicate. After genomic DNA was extracted from the library, the transposon adjacent regions were enriched by PCR before massive parallel sequencing. The resultant sequencing data were analyzed using TRANSIT software (26). TA dinucleotides of all libraries were adequately saturated (>30% saturation) and were included in the further analysis (Table S3). Gene essentiality was assessed using the Hidden Markov Model (HMM) method by TRANSIT. We identified 2744 to 3495 non-essential genes, 121 to 224 essential genes, 120 to 576 growth-advantage genes, and 124 to 220 growth-defect genes (Figure 3A, Table S4, S5, and S6). All plasmid-encoded genes were predominantly non-essential except for several growth-defect genes and growth-advantage genes (Figure S1, Table S4, S5, and S6).

**Figure 3.**
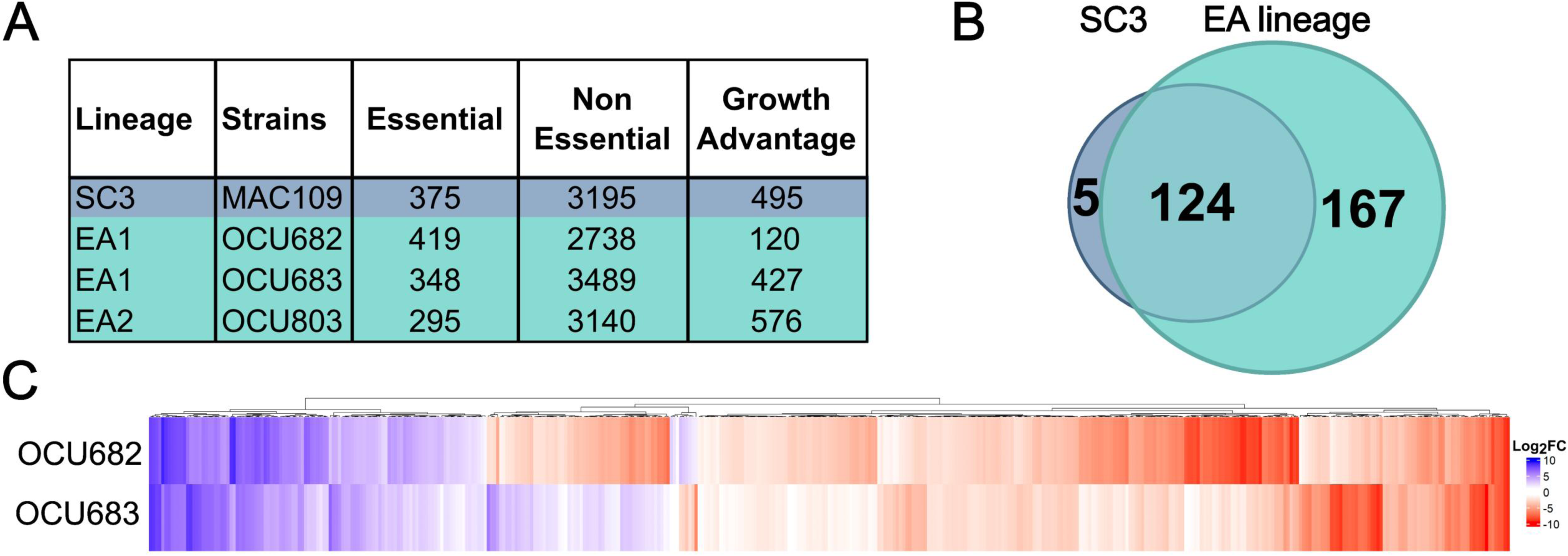
Gene classification identified by TRANSIT and gene comparison of lineage and among EA lineage strains. A) *In vitro* classification of genes in OCU682, OCU683, OCU803, and MAC109 using TRANSIT. The numbers of genes classified as essential (a category that includes both essential and growth-defect), non-essential, and growth-advantage, after excluding low-confidence genes filtered out by TRANSIT HMM analysis, are shown. B) Venn diagram showing the *in vitro* essential genes in EA lineages, which included OCU682, OCU683, OCU803, and SC3 lineage, i.e., MAC109, excluding low-confidence genes filtered out by TRANSIT HMM analysis. Each Venn diagram color is identical to Figure 3A. C) Gene essentiality profiles of OCU682 and OCU683 were compared to OCU803 by a resampling-based method in TRANSIT. The heatmap displays the relative essentiality differences across strains, with genes hierarchically clustered based on their essentiality profiles, taking into account statistical significance (adjusted p-value < 0.05). The left dendrogram indicates similarity in essentiality patterns among genes. Only genes with adequate transposon insertion coverage in all strains were included in the analysis.

### Reanalysis of MAC109 for standardized comparison

To enable direct comparison with the EA lineage datasets, MAC109 was reanalyzed with the same pipeline to eliminate methodological biases (22). Using this uniform pipeline, MAC109 contained 3,195 non-essential genes, 216 essential genes, 495 growth-advantage genes, and 159 growth-defect genes (Figure 3A, Table S7). Most plasmid-encoded genes were categorized as non-essential, with the exception of four growth-defect genes and eighteen growth-advantage genes (Figure S1, Table S7).

### Comparative analysis across lineages

Across the three EA strains and MAC109, we identified 124 pan-essential genes (Figure 3B, Table S8). These included functions related to DNA replication and repair, protein translation, and central metabolic pathways. Several known antitubercular drug targets (*inhA, embAB, gyrAB, rpoB, mmpl3,* and *dprE1*) were also pan-essential. Interestingly, among the eight penicillin-binding proteins (PBPs) in MAH, only *pbpB* was consistently essential, while the remaining homologs were dispensable (Table S8, Figure S3). We further identified 5 genes uniquely essential in MAC109 and 167 uniquely essential in EA lineages. Most EA-specific essential genes encoded proteins related to lipid metabolism and cell envelope biosynthesis (e.g., acyl-CoA dehydrogenases, polyketide synthases, O-antigen ligases). In addition, multiple type VII secretion system components, PE/PPE family proteins, and Mce transporters were essential exclusively in EA strains. Notably, most of these EA-specific essential genes belonged to the core genome but were non-essential in MAC109.

To assess differences in gene essentialities within EA lineages, we applied the resampling method in TRANSIT to compare EA1 and EA2 strains with the EA2 reference OCU803.

Among 3,777 orthologous core genes, 439 showed statistically significant differences in essentiality (Figure 3C, Table S9). These genes were enriched in central metabolism, redox homeostasis, and cell envelope biosynthesis, suggesting fine-scale, lineage-dependent adaptation of survival pathways.

## Discussion

Despite the rising global incidence of MAH-related pulmonary disease, our understanding of MAH gene essentiality remains limited. MAH exhibits extensive genomic diversity, yet the functional implications of this diversity and the conservation of essential genes across lineages remain poorly understood. In this study, we combined Tn-Seq and whole-genome sequencing of three EA isolates with published SC3 data to perform a comparative functional genomic analysis. This dual approach enabled us to identify both core-essential genes and lineage-specific essential genes.

The pan-essential genes identified in both EA and SC lineages highlight fundamental processes indispensable for MAH survival, including DNA replication and repair, protein translation, and cell envelope biogenesis. Representative examples include genes encoding ribosomal subunits, aminoacyl-tRNA ligases, arabinosyltransferases, as well as *fadD32* and *embB*, underscoring the central roles of genetic information processing, protein synthesis, and the lipid-rich cell envelope. Conserved chaperones and proteases such as GroEL and Clp further emphasize the importance of proteostasis and stress tolerance. Together, these genes define a conserved genetic backbone upon which lineage-specific adaptations are superimposed.

In addition to defining a universal set of core-essential genes, our study revealed that the EA lineage-specific essential genes were enriched in the genes involved in cell envelope remodeling, secretion systems, and redox balance. These pathways are critical for host adaptation and may contribute to the higher prevalence and drug resistance frequently observed in the EA lineage (27). For example, the essentiality of multiple *mce* transporters and type VII secretion components in EA isolates suggests selective pressures to withstand host-associated stresses (28, 29). Similarly, the unique requirement for lipid metabolism genes underscores the metabolic flexibility of these lineages in nutrient-limited environments.

We also found differences in gene requirements between EA1 and EA2 strains. This observation indicates that essentiality can diverge not only between EA and SC, but also within closely related lineages. Such lineage-dependent resiliency has been reported not only in *Mtb* but also in *Streptococcus pneumoniae*, where it contributes to clinically relevant phenotypes (30). Considering that each MAH strain harbored more than 500 accessory genes, our results underscore the plasticity of essentiality and its modulation by the accessory genome, consistent with recent pan-genome studies in other pathogens (16, 31, 32). This plasticity poses both challenges and opportunities for drug development. From a therapeutic perspective, the limited set of pan-essential genes (e.g., *pbpB*, *dprE1*, *inhA*) emphasizes the challenge of finding universally conserved targets. In particular, penicillin-binding proteins (PBPs), the targets of β-lactams, were largely non-essential, with *pbpB* being the only consistently essential homolog across all strains. Although this study analyzed a relatively small number of strains and further experimental validation will be required, these findings suggest physiological adaptations that may influence drug susceptibility. Taken together, our results highlight both the species-specific patterns of essentiality and the need to distinguish conserved from lineage-variable targets when prioritizing therapeutic strategies.

## Material and Methods

### Bacterial strains, media, and growth conditions

MAH strains, OCU682, OCU683, and OCU803, were isolated from a patient’s residential bathrooms (33). MAH strains were grown aerobically at 37°C in Middlebrook 7H9 (BD Difco^TM^) medium supplemented with oleate-albumin-dextrose-catalase (OADC; 10%, vol/vol), glycerol (0.2%, vol/vol), and tyloxapol (0.05%, vol/vol).

### Whole genome sequencing by PacBio system

Genomic DNA was extracted from the cell pellet derived from cultured medium using the conventional phenol–chloroform method after bead beating (0.1 mm zirconia beads; Vortex Mixer GENIE2 with Microtube Attachment at max speed for 7 min) as described previously (34). The concentrations of DNA were measured by QuantiFluor dsDNA System on Quantus Fluorometer (Promega). The DNA quality was confirmed by Agilent HS Genomic DNA 50 kb Kit on 5200 Fragment Analyzer System (Agilent Technologies).

Single-Molecule Real Time (SMRT) sequencing service provided by the Bioengineering Lab. Co., Ltd (Saitama, Japan) was used. Genomic DNA was purified by using DNA Clean Beads (MGI Tech Co., Ltd.) and fragmented to approximately 10–20 kbp using ME220

Focused-ultrasonicator (Covaris). The DNA sequencing library was prepared using the SMRTbell Express Template Prep Kit 2.0 (PacBio) following the Procedure & Checklist instructions. Polymerase complexes were prepared by Revio Polymerase kit (PacBio).

Sequencing was conducted using Pacific Biosciences Revio. Among the three MAH strains, a complete genome of OCU683 was successfully assembled using PacBio Revio data.

### Additional whole genome sequencing using Illumina and Pacbio systems

For OCU682 and OCU803, additional sequencing was required to obtain complete genome assemblies. Strains were inoculated into Middlebrook 7H11 agar (BD Difco^TM^) supplemented with 10% Middlebrook OADC and incubated at 37°C for 2-3 weeks. DNA was extracted from bacteria clumps in one volume of inoculation loops (4 mm) using the conventional phenol–chloroform method after bead beating (0.2 mm glass beads; Vortex Mixer GENIE2 with Microtube Attachment at max speed for 5 min).To complete genome assemblies, OCU682 was supplemented with Illumina NovaSeq 6000 paired-end reads, while OCU803 was supplemented with both Illumina NovaSeq 6000 and additional long-read data from the PacBio Sequel II platform. These supplementary sequencing efforts were conducted at the University of Minnesota Genomics Center (Minneapolis, MN, USA).

### Whole genome sequencing analysis

SMRT Link (ver. 12.0.0.177059) (PacBio) was used to remove overhanging sequence adaptors, and consensus sequence reads with an average quality value of less than 20 per read were removed. Filtlong (version 0.2.1) (https://github.com/rrwick/Filtlong) was used to eliminate reads shorter than 1,000 bases. *De novo* assembly was performed using Flye (ver. 2.9.2-b1786) (35) and Bandage (ver. 0.8.1) (36). The completeness of the genome was assessed using CheckM2 (ver. 1.0.1) (37). The complete sequence of the OCU683 strain was obtained by the analysis above.

To obtain the complete sequences of OCU682 and OCU803, we conducted hybrid *de novo* genome assembly by incorporating additional Illumina sequencing data. For OCU803, additional long-read data from the PacBio Sequel II platform were also used. Hybrid assemblies were generated using Flye (ver. 2.9.2-b1801) and Unicycler (ver. 0.5.1) (38). Genome annotation for all three strains was performed using the NCBI Prokaryotic Genome Annotation Pipeline (PGAP) with default settings (39).

### Lineage classification

The 185 genome sequences used for lineage classification are listed in Table S2. Genome data for 182 strains were retrieved from the NCBI RefSeq database for entries registered as “*Mycobacterium avium* subsp. *hominissuis”* (1st May 2025). Core genome SNPs were extracted using the parsnp v.2.1.1 (40), with the genome sequence of strain TH135 (41) as the reference, as described previously (7–9). SNPs located in locally collinear blocks (LCBs) smaller than 200 bp, as well as sites containing gaps or ambiguous bases (N), were removed using harvesttools (42). To determine optimal population clusters, the haplotype information of the filtered SNPs was analyzed using a Bayesian hierarchical clustering algorithm in fastBAPS (43). Visualization of the phylogenetic tree was done using iTOL (44).

### Construction of saturated transposon libraries of MAH

Saturated transposon libraries were constructed essentially as previously described (17). The protocol was optimized by using pre-warmed both buffer and phage, and maintaining continuous shaking during phage transfection for no more than 20 hours. Mycobacteriophage phAE180 (45), at a titer of 2.12 to 11.0 ×10^10^ pfu/ml, was used to transduce a mariner derivative transposon Tn5371 (46) into three strains grown to an OD_600_ of 0.4 to 0.9. The transduced cells were resuspended in 7H9 medium and spread on a 7H9 agar plate containing tyloxapol (0.05%, vol/vol) and 50 µg/ml kanamycin, then incubated at 37 °C for 7–14 days. The resulting mutant libraries were scraped off the plates and stored at -80°C for further genomic DNA extraction. Each transposon library was generated in duplicate.

### Transposon sequencing (Tn-Seq)

The genomic DNA (gDNA) was extracted as described in the above section. Tn-Seq sequencing library preparation was performed essentially as previously described (47). Briefly, 2 mM MgSO_4_ solution containing the oligonucleotides, 5′-TACCACGACCA-NH_2_ and 5′-ATGATGGCCGGTGGATTTGTGNNA NNANNNTGGTCGTGG TAT, each at 100 μM, was heated at 95°C for 10 minutes and gradually cooled to 20°C to prepare the barcoded adaptor. gDNA was fragmented using the ME220 Focused-ultrasonicator (Covaris).

End-repairing and ligation of the barcoded adaptors to the fragmented gDNA were performed using an NEBNext Ultra II DNA Library Prep Kit for Illumina (New England Biolabs) according to the manufacturer’s instructions. The barcoded adaptor ligated gDNA fragments were purified with the QIAquick PCR purification kit (QIAGEN). Transposon junctions were amplified by using the transposon specific primer T7 and the adapter specific primer JEL_API with GoTaq colorless Master Mix (Promega) supplemented with 5% (vol/vol) DMSO with the following PCR condition (95°C for 10 min, 20 cycles of 95°C for 30 seconds, 58°C for 30 seconds, and 72°C for 45 seconds, and final extension at 72°C for 5 minutes). Amplification products were purified with Sera-Mag Select (Cytiva). Adapter sequences for Illumina sequencing were added using the two-step PCR reactions below. The first PCR was performed using the Pre-Index primer mixture and Pre-Universal primer mixture with NEBNext Ultra II Q5 master mix with the following PCR condition (98°C for 10 min, 10 cycles of 98°C for 10 seconds, 70°C for 30 seconds, and 72°C for 30 seconds, and final extension at 72°C for 2 minutes). Index PCR was performed using NEBNext Multiplex Oligos. The resultant Tn-Seq library was sequenced using a NextSeq 2000 (Illumina), 125 bp PE run using NextSeq 1000/2000 P1 XLEAP-SBS Reagents (Illumina). All primers used in this study are listed in Table S10.

### Tn-Seq analysis

Tn-Seq data were processed using the TPP tool from the TRANSIT v3.3.4 analysis platform, and transposon genome junctions were mapped to the assembled genomes of the isolate using the Burroughs-Wheeler aligner (BWA) (26). Tn-Seq data of MAC109 from NCBI’s BioProject database under accession number: PRJNA527645 (22). MAC109 genome sequence was obtained from NCBI RefSeq and reannotated by the NCBI PGAP. This was necessary in order to perform the analysis with identical procedures and parameters, thereby minimizing methodological biases and ensuring that observed differences reflected true biological variation.

Gene essentiality was assessed using the Hidden Markov Model (HMM) method by TRANSIT. The HMM method is based on the read count at a given site and the distribution over the surrounding sites. After the initial gene calls using the HMM method, HMM confidence scores were calculated based on the consistency of the insertion statistics with the posterior distribution of the essentiality state. Low-confidence gene calls below 0.20 were excluded. Mutual orthologues among OCU682, OCU683, OCU803, and MAC109 were defined by reciprocal best-hit analysis using BLASTp with an e-value cutoff of 1×10⁻^10^. Strain- or lineage-specific essentiality was assessed using the resampling method implemented in TRANSIT, relative to OCU803. The *p*-values were adjusted for multiple comparisons using the Benjamini–Hochberg procedure with a false discovery rate threshold of 5%. Genes with adjusted *p*-values (*p*-adj) < 0.05 were considered significantly different in essentiality.

### Pan-genome analysis

For the pan-genome analysis, 26 complete genomes were used (Table S2). A pairwise mash distance was calculated using Mashtree v.1.4.6 (48). All genome sequences were reannotated by the NCBI PGAP. The pan-genome analysis was conducted using Roary v.3.11.2 with default parameters and identified the core, soft-core, shell, and cloud genes (49).

## Data availability

All genome sequences have been deposited in GenBank under accession numbers AP042353 to AP042360 and in the NCBI BioProject database under accession number PRJDB35554.

## Acknowledgements

This study was financially supported by funds from the Japan Agency for Medical Research and Development (JP23wm0325037 to Y.M., JP25fk0108673 to F.M.), JSPS KAKENHI (23H02634 and 23K27325 to Y.M., 24K1163 to Y.N., and 23H00451 to F.M.).

